# Allele-specific open chromatin analysis revealed allelic bias of alcohol-induced gene expression in human microglia

**DOI:** 10.64898/2026.11.23.768544

**Authors:** Xindi Li, Nicholas C. Green, Jaime S. Messenger, Shreeya Govande, Henri D Noel, Tanya Soumine, Yue Wang, Jubao Duan, Yunlong Liu, Dongbing Lai, Ronald P. Hart, Zhiping P. Pang

**Author notes:** Co-corresponding authors Dongbing Lai, Ronald P. Hart, Zhiping P. Pang.

## Abstract

Alcohol use disorder (AUD) is a polygenic disorder. Most AUD-associated variants identified by genome-wide association studies (GWAS) are located in non-coding regions, making it challenging to determine which variants are functional and how they contribute to disease-relevant cellular phenotypes. Here, we integrated microglia-specific allele-specific open chromatin (ASoC) mapping, GWAS, eQTLs, differentially expressed genes identified from postmortem human microglia and iPSC-derived microglial-like cells, and chromatin accessibility (ATAC-seq) data from postmortem microglia to prioritize regulatory variants, followed by functional validation using multiple assays. We identified a panel of AUD-associated non-coding variants, particularly rs7307261, as a candidate functional variant that showed regulatory potential. The minor allele (A) of rs7307261 showed higher predicted similarity to binding motifs for E26 transformation-specific (ETS)-family transcription factors, including FLI1 and ELF2, than the G allele. The A allele was also associated with increased *CD163* expression in AUD postmortem microglia and in iPSC-derived microglial-like cells after ethanol exposure. Notably, the ethanol-induced *CD163* expression was abolished by knocking out rs7307261 or converting rs7307261 A/A to G/G, accompanied by changes in ethanol-induced microglial phagocytic function. Together, our study provides a framework for translating AUD genetic associations into context-specific functional mechanisms by linking non-coding risk variants to target genes and ethanol-induced cellular phenotypes.

## Introduction

Alcohol use disorder (AUD) is a chronic and relapsing disorder due to both environmental exposures and genetic predisposition. Recent large-scale genome-wide association studies (GWAS) have identified hundreds of variants associated with AUD, highlighting the strong genetic contribution to this disorder^1, 2^. However, most AUD-associated variants lie in non-coding regions, and many may simply tag truly functional variants in linkage disequilibrium, making it challenging to identify those that have functional impact. Furthermore, how functional variants contribute to the disease risk remains to be discovered.

Open chromatin regions (OCRs) affect gene expression^3^, and allele-specific open chromatin (ASoC) mapping provides an effective approach to identify non-coding variants with regulatory function^4–7^. ASoC compares chromatin accessibility between two alleles of heterozygous variants within the same individual, and allelic imbalance indicates potential allele-specific regulatory effects in disease-relevant cell types. In human iPSC-derived neurons and microglia, ASoC has been used to identify functional non-coding variants in schizophrenia^4, 5^ and Alzheimer’s disease^3^, respectively. Importantly, studies in both human and animal models have shown that chronic alcohol consumption induced widespread changes in chromatin accessibility across the brain, suggesting that AUD-associated variants located within OCRs may regulate gene expression in the context of ethanol exposure^8–11^. Therefore, by integrating cell-type-specific ASoC mapping with AUD-related expression quantitative trait loci (eQTLs) and differentially expressed genes (DEGs) between AUD cases and controls, we can prioritize AUD-associated variants with regulatory potential and identify their target genes.

Recent research has increasingly highlighted the impact of chronic alcohol consumption on neuroimmune regulation, particularly within the central nervous system^12, 13^, where alcohol activates microglia and alters neuroimmune signaling pathways, potentially contributing to AUD. Microglia, the brain’s resident immune cells, are of particular interest given the strong evidence for immune and inflammatory pathway activation in AUD^14–17^. Additionally, gene expression profiling of postmortem human brains has revealed distinct microglial activation states in AUD^11^. Consistent with this, our previous work demonstrated that chronic alcohol consumption altered microglial major histocompatibility complex (MHC) class II expression and impaired phagocytic function^17^. Therefore, microglia are a key cellular context for understanding the functional impact of AUD-associated variants on shaping the neuroimmune responses associated with chronic alcohol consumption.

In this study, we aimed to identify functional non-coding AUD variants and their target genes by systematically integrating microglia-specific ASoC mapping, GWAS^2^, eQTLs, and DEGs between AUD cases and controls from both postmortem microglia and human microglia cultures exposed to ethanol^11, 17^. Rather than relying solely on overlapping associations, we ensured that each association’s direction remained consistent across all analytical steps, and that gene expression changes matched predictions from allele-specific functions. We identified a panel of ASoC variants associated with AUD. We found that carriers of the A allele of the non-coding variant rs7307261 had increased AUD risk, driven by increased *CD163* expression at both RNA and protein levels. Functional assays and CRISPR editing demonstrated its potential link to microglial phagocytic function in the context of ethanol exposure. Collectively, our work outlines a multiscale framework that connects AUD risk variants from the genetic level, through molecular regulation of gene expression, to downstream cellular changes, providing a path from genetic association to potential disease mechanisms.

## Results

To identify functional variants associated with AUD, we used an integrated and streamlined workflow. First, we identified AUD-associated variants from AUD-related GWAS that were located in microglia-specific ASoC^3^. Next, we prioritized variants that were also identified as eQTLs^11^. We then used the same human postmortem microglia RNA expression dataset^11^ to prioritize candidate target genes differentially expressed between AUD cases and controls and further confirmed these genes in human microglial-like cell cultures^17^. We applied a directional consistency filter to select variant-gene pairs showing concordant relationships among allele-specific chromatin accessibility, gene expression patterns, and AUD risk associations. We confirmed the directions of allele-specific disease risk and chromatin status with ATAC-seq data^11^. Lastly, we validated the variant and its target gene’s role in AUD using multiple assays and CRISPR editing.

### Microglial ASoC prioritizes AUD GWAS variants with regulatory potential

To prioritize AUD-associated variants with potential regulatory effects in microglia, we integrated AUD-related GWAS results with microglia ASoC regions (**Fig. 1a)**. Given the highly polygenic nature of AUD, we included all GWAS variants nominally associated with AUD (P < 0.05) to capture all potential regulatory variants. Of these, 67,101 variants mapped to microglial ASoC regions, including 1,316 identified as eQTLs in the human postmortem microglial gene expression dataset^11^ (**Fig. 1b, Supplementary Fig. 1a, Supplementary Table 1**). ASoC-overlapping AUD-associated SNPs showed significant enrichment across multiple genomic and functional annotations, particularly in intronic and upstream/downstream regions, with the strongest enrichment observed for eQTLs (**Fig. 1c**). After aligning directions of effect of eQTLs and AUD-associated variants, 366 variants were associated with decreased target gene expression and lower AUD risk, 308 variants with increased target gene expression and lower AUD risk, 242 variants with decreased target gene expression and higher AUD risk, and 400 variants with increased target gene expression and higher AUD risk (**Fig. 1d, Supplementary Table 2**).

**Figure 1:**
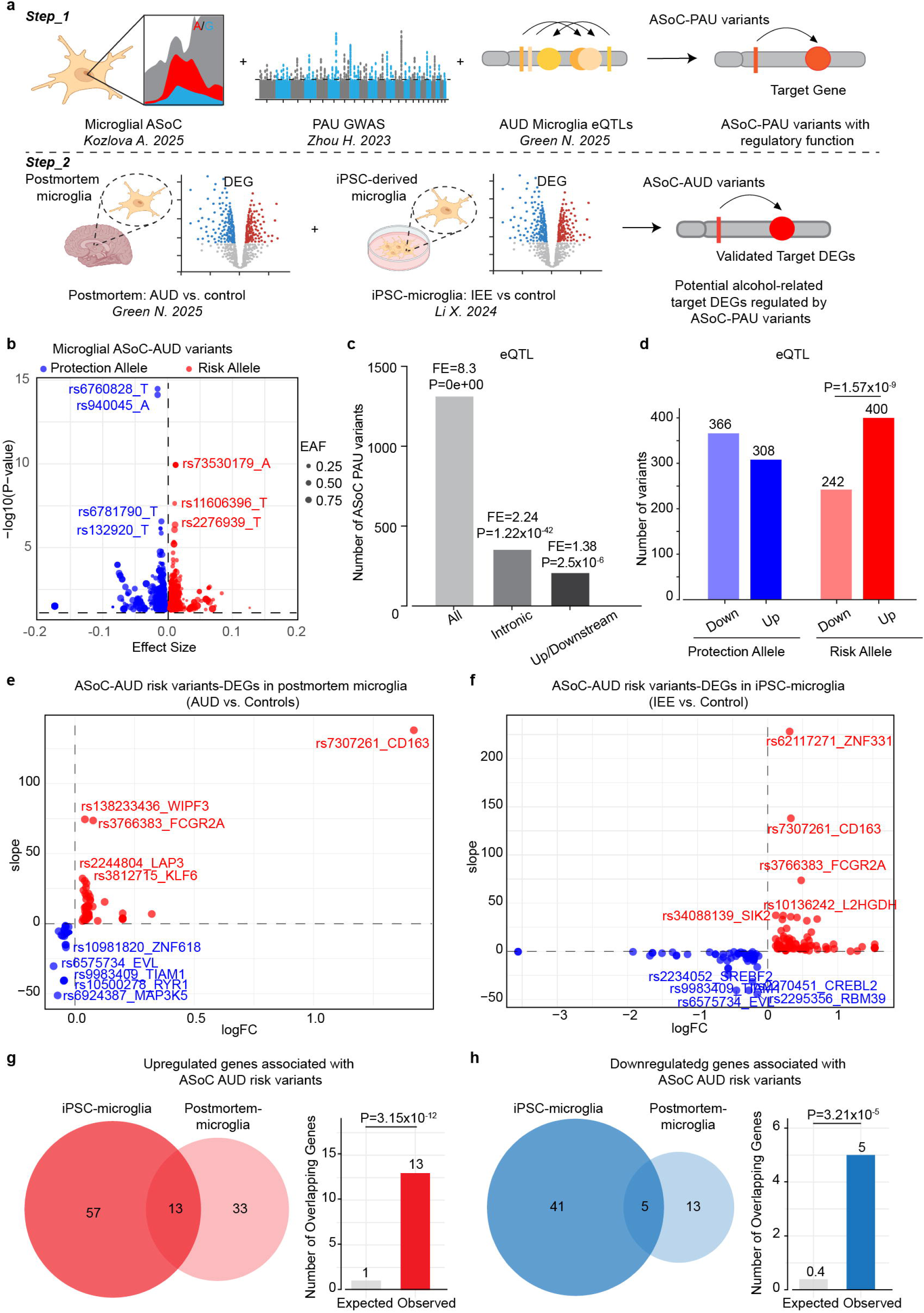
Identification of AUD-associated microglial ASoC variants. **(a)** Workflow for prioritizing functional AUD-associated variants in microglia by integrating microglial ASoC variants, AUD-related GWAS, and AUD microglia eQTLs, and identifying candidate ASoC-AUD variant target genes using DEGs from postmortem microglia from AUD cases and controls and ethanol-exposed iPSC-derived microglia. **(b)** Volcano plot of microglial ASoC-AUD variants, where red and blue indicate increased and decreased AUD risk, respectively. **(c)** Enrichment of ASoC variants relative to all AUD GWAS variants with P < 0.05 across regulatory annotations, including variants with eQTL evidence, intronic variants, and variants located in upstream or downstream regulatory regions. FE indicates fold enrichment. P values were calculated using a hypergeometric test to assess enrichment of ASoC-AUD variants in each annotation category compared with the background variant set. P <0.05 was considered significant. **(d)** Bar graph showing the number of protective and risk AUD variants associated with upregulated (red) or downregulated (blue) target genes. P values were calculated using a chi-square test. P <0.05 was considered significant. **(e, f)** Volcano plots showing DEGs associated with AUD-ASoC risk alleles in AUD postmortem brain (b) and microglial cultures (c). Red indicates genes upregulated by risk alleles, and blue indicates genes downregulated by risk alleles. **(g)** Venn diagram showing the overlap of upregulated DEGs associated with ASoC-AUD risk alleles between microglial cultures and postmortem microglia. Bar graphs show enrichment of overlapping DEGs relative to all DEGs. P values were calculated using a one-sided hypergeometric test, with P < 0.05 considered significant. **(h)** Venn diagram showing the overlap of downregulated DEGs associated with ASoC-AUD risk alleles between microglial cultures and postmortem microglia. Bar graphs show enrichment of overlapping DEGs relative to all DEGs. Bar graphs show enrichment of overlapping DEGs relative to all DEGs. P values were calculated using a one-sided hypergeometric test, with P < 0.05 considered significant.

### Potential target genes regulated by microglial ASoC prioritized AUD-associated variants

To prioritize potentially biologically relevant variant-gene pairs, we first identified DEGs in human postmortem microglia from AUD cases versus controls whose expression changes were consistent with directions of effect of eQTLs. We then examined whether these genes also showed concordant differential expression in our previous human microglial-like cell cultures with and without ethanol exposure **(Fig. 1a).**

In total, we identified 74 variant-gene pairs (64 DEGs) in which regulatory directions of the effective allele matched the gene expression changes observed in microglia from AUD cases versus controls **(Fig. 1e, Supplementary Table 3),** including 46 upregulated and 18 downregulated genes. We also identified 139 variant-gene pairs (116 DEGs) in which regulatory directions of the effective allele matched the gene expression changes observed after ethanol exposure in cultured human microglial-like cells **(Fig. 1f, Supplementary Table 3),** including 70 upregulated and 46 downregulated genes. Across the two datasets, 13 genes in 19 variant-gene pairs were upregulated in both, including *CD163* and *FCGR2A* (**Fig. 1g, Supplementary Fig. 1b**), while 5 genes, including *TIAM1* and *EVL*, in 6 variant-gene pairs were downregulated in both (**Fig. 1h, Supplementary Fig. 1c**). This number of overlaps was significantly greater than expected by chance (P = 3.15x10^-12^ for upregulated genes, P = 3.21x10^-5^ for downregulated genes; **Fig. 1g, h**), supporting the prioritization of these variant-gene pairs for further investigation.

### The rs7307261 A allele is associated with increased chromatin accessibility in AUD microglia

Having prioritized these ASoC variant-gene pairs, we next asked whether chromatin regions containing these variants showed differential accessibility between AUD cases and controls. To address this, we analyzed ATAC-seq data from human postmortem microglia from AUD cases and controls **(Fig. 2a)**. Across all variant-gene pairs examined, the region surrounding rs7307261 showed a significant increase in chromatin accessibility in AUD microglia compared with controls (**Supplementary Fig. 2a, b**), suggesting that chronic alcohol consumption may alter the open chromatin state at this locus. To assess genotype-specific effects, we separated microglia by genotype at rs7307261 (i.e., A/A, G/A, and G/G). We found that the rs7307261 A/A genotype showed significantly higher accessibility around rs7307261 in AUD cases compared with controls (**Fig. 2b**), while the G/A (**Fig. 2c**) and G/G (**Fig. 2d**) genotypes showed no significant changes. When comparing allele-specific accessibility directly, the rs7307261 A allele showed a significant increase of chromatin accessibility in AUD microglia, whereas the G allele did not (**Fig. 2e**). These findings indicate greater chromatin accessibility associated with the A allele due to chronic alcohol exposure.

**Figure 2:**
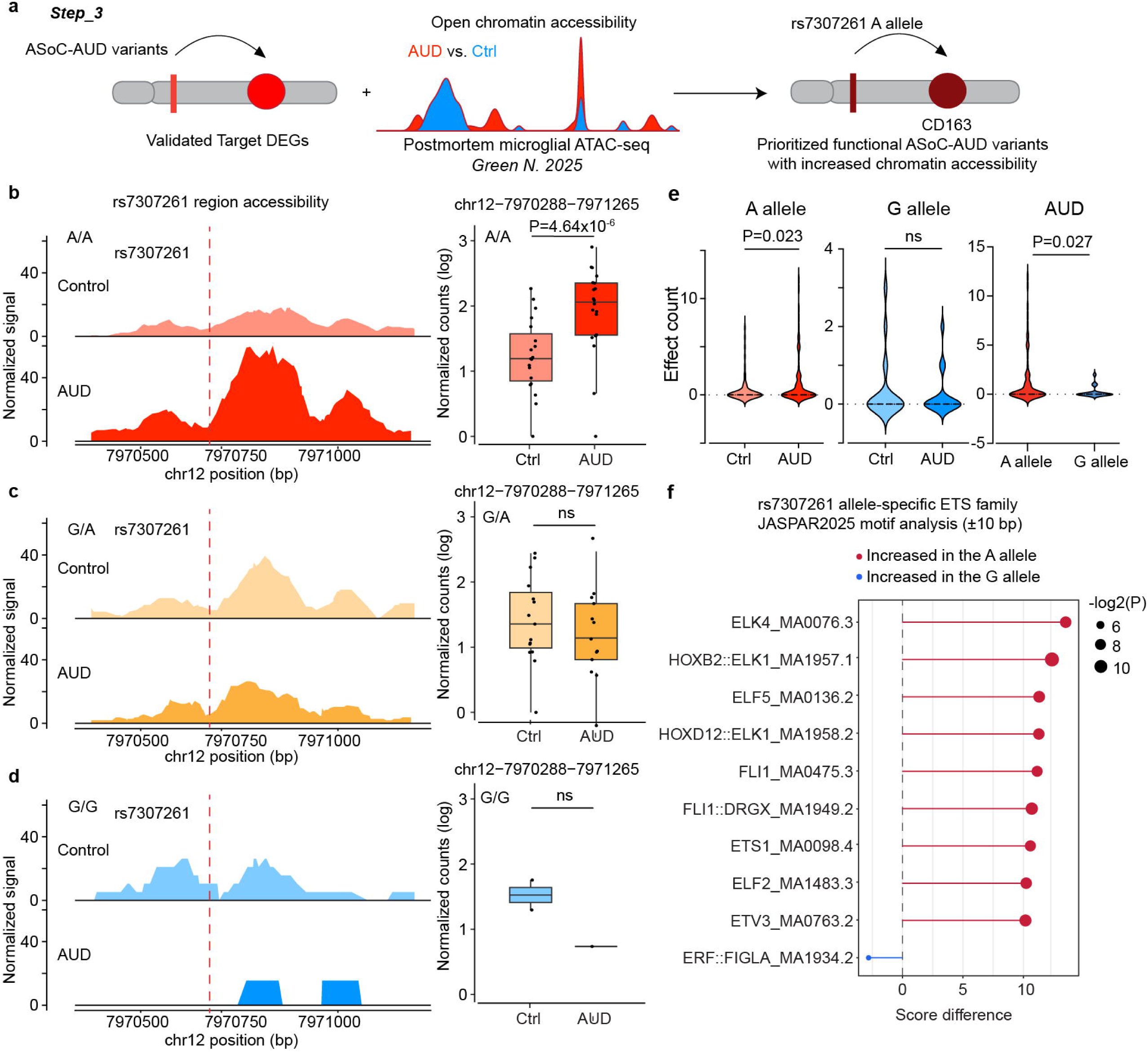
rs7307261 A allele is associated with increased chromatin accessibility in human microglia. **(a)** Workflow of integrating validated ASoC-AUD variant target genes with postmortem microglial ATAC-seq data to prioritize functional variants showing increased chromatin accessibility in human postmortem microglia. **(b-d)** Chromatin accessibility at the rs7307261 region across rs7307261 genotypes in AUD and control postmortem microglia. Accessibility profiles are shown for A/A (b, red), G/A (c, yellow), and G/G (d, blue) microglia from control and AUD cases. Quantification of chromatin accessibility at the rs7307261 locus in microglia from AUD cases and controls stratified by genotype: A/A (c; AUD, n = 20; control, n = 19), G/A (e; AUD, n = 13; control, n = 15) and G/G (g; AUD, n = 1; control, n = 2). ns, not significant. **(e)** Violin plots of chromatin accessibility effect count at the rs7307261 locus. The left plot shows accessibility at rs7307261 for the A allele in control and AUD microglia. The middle plot shows accessibility at rs7307261 in control and AUD microglia. The right panel compares accessibility at rs7307261 between the A and G alleles in AUD microglia. Statistical analysis was performed using a two-tailed Mann-Whitney test. Data are presented as mean ± SEM; P < 0.05; ns, not significant. **(f)** JASPAR 2026 motif analysis comparing predicted ETS family transcription factor binding scores between the A and G alleles using ±10 bp sequence surrounding rs7307261. Red indicates higher motif scores for the A allele, whereas blue indicates higher scores for the G allele. Dot size represents the -log₂-transformed P value.

To test whether this allele-specific accessibility reflects altered transcription factor binding potential, we analyzed the sequence surrounding rs7307261 using the JASPAR 2026 motif database. The rs7307261 A allele showed higher predicted similarity to binding motifs for E26 transformation-specific (ETS)-family transcription factors, including FLI1 and ELF2, than the G allele (**Fig. 2f, Supplementary Fig. 2c**). These factors have been implicated in microglial development and the regulation of immune-related transcriptional programs^18^. Together, these results suggest that chronic alcohol consumption preferentially increases chromatin accessibility for carriers of the rs7307261 A allele, potentially enhancing a microglia-relevant regulatory motif at this locus.

### The rs7307261 A allele is associated with upregulation of CD163 in microglia following ethanol exposure

Both eQTL and DEG analyses indicated that *CD163* was a potential target gene of rs7307261. To further investigate the relationship between the rs7307261 A allele and *CD163* expression, we compared *CD163* mRNA levels across rs7307261 genotypes in postmortem microglia and iPSC-derived microglial-like cells (A/A, n = 11 cell lines; G/A, n = 9; G/G, n = 4). *CD163* mRNA levels were higher in A/A than in G/A microglia in both postmortem microglia and human iPSC-derived microglial-like cells; however, the difference was no longer significant after adjusting for AUD status in postmortem microglia **(Fig. 3a)**. In addition, *CD163* expression was significantly higher in A/A than in G/G iPSC-derived microglial-like cells, whereas this difference was not significant in postmortem microglia. These findings showed that the association between rs7307261 genotype and *CD163* expression is not consistently observed in the absence of chronic alcohol consumption or ethanol exposure, suggesting that the regulatory effect of the A allele may be alcohol dependent.

**Figure 3:**
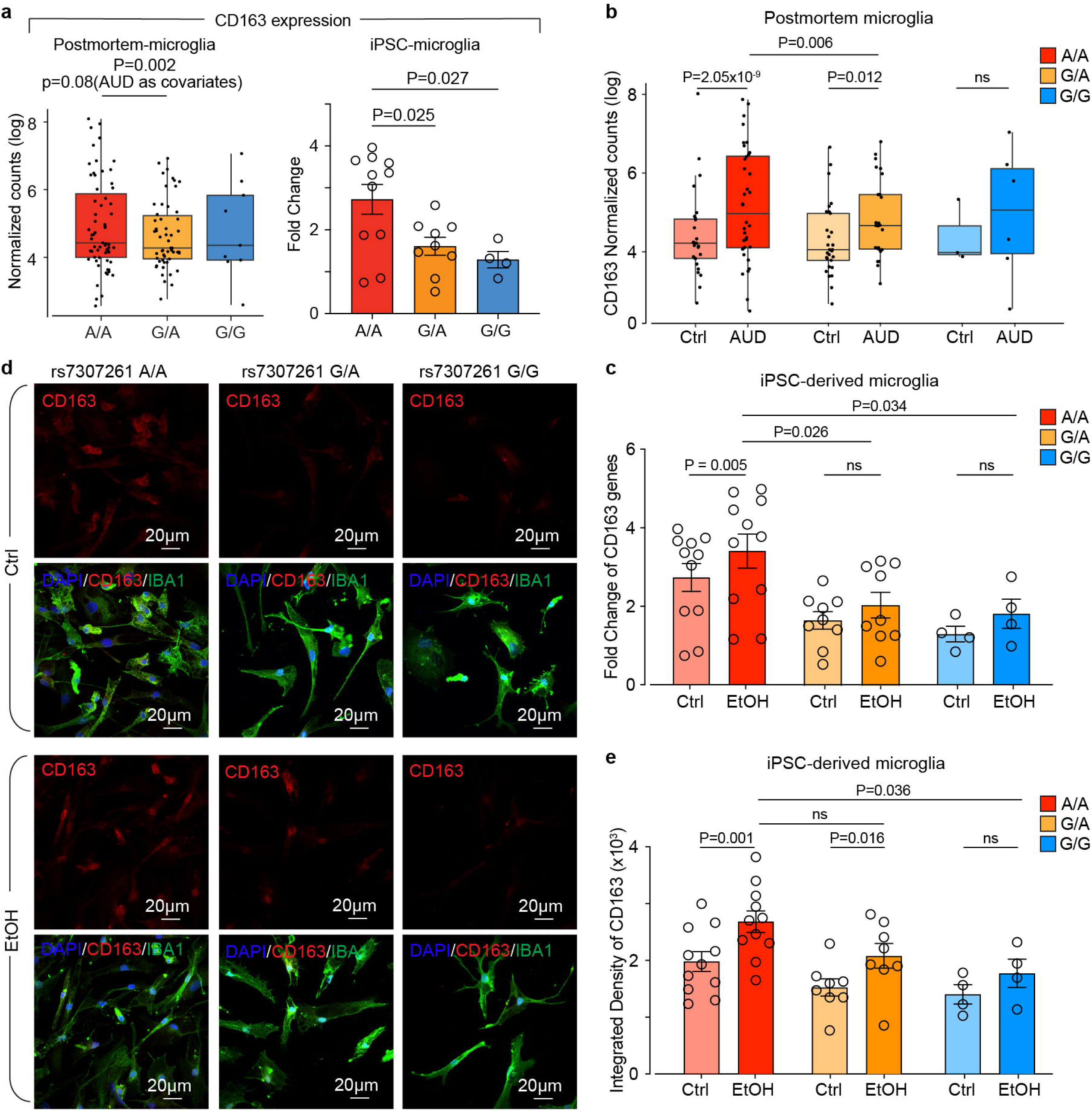
rs7307261 A allele is associated with increased *CD163* expression in AUD postmortem microglia and iPSC-derived microglia cells after ethanol exposure. **(a)** Comparison of CD163 expression across rs7307261 genotypes (A/A=63, G/A=54, G/G=9) in postmortem microglia from individuals with AUD and iPSC-derived microglial-like cells (A/A=11, G/A=9 and G/G=4). **(b)** Bar graph showing *CD163* mRNA expression across rs7307261 genotypes in postmortem microglia from AUD cases and controls: A/A (AUD, n = 36; control, n = 27), G/A (AUD, n = 25; control, n = 29) and G/G (AUD, n = 6; control, n = 3). **(c)** Bar graph showing *CD163* mRNA expression measured by RT-qPCR in iPSC-derived microglial cells with or without ethanol exposure across rs7307261 genotypes: A/A (n = 11 cell lines), G/A (n = 9 cell lines), and G/G (n = 4 cell lines). Paired data were analyzed using a mixed-effect model with the Geisser-Greenhouse correction, followed by Šídák’s multiple-comparisons test. Data are presented as mean ± SEM. NS, not significant. **(d)** Representative immunofluorescence images showing CD163 (red), IBA1 (green), and DAPI (blue) staining in human microglial cells carrying rs7307261_A/A, G/A, and G/G genotypes with or without ethanol exposure. **(e)** Bar graph showing quantification of CD163 immunostaining in iPSC-derived microglial cells across rs7307261 genotypes with or without ethanol exposure: A/A (n = 11 cell lines), G/A (n = 8 cell lines) and G/G (n = 4 cell lines). Paired data were analyzed using a mixed-effect model with the Geisser-Greenhouse correction, followed by Šídák’s multiple-comparisons test.

As the rs7307261 A allele carriers showed increased chromatin accessibility in AUD cases, we next tested whether the rs7307261 A allele further altered *CD163* expression levels under alcohol-related conditions. In rs7307261 A/A microglia, *CD163* expression increased in both AUD postmortem samples and human microglial-like cells following ethanol exposure **(Fig. 3b, c).** G/A microglia also showed increased *CD163* mRNA expression in AUD postmortem samples, although this increase was not significant in iPSC-derived microglial-like cells (**Fig. 3b, c**). In contrast, G/G microglia showed no significant change in *CD163* mRNA expression in either dataset (**Fig. 3b, c**). We further validated these findings at the protein level by immunostaining (**Fig. 3d**). Consistent with the mRNA results, ethanol increased CD163 fluorescence intensity in A/A and G/A microglial-like cells, but not in G/G lines, with a greater increase in A/A than in G/G cells (**Fig. 3e**). Together, these results suggest that the rs7307261 A allele was associated with increased *CD163* gene and protein expression under ethanol exposure.

To further test the allele-specific effect of rs7307261 within the same genetic background, we used CRISPR editing in an rs7307261 A/A iPSC line. We generated a knockout line by deleting a 17bp/18bp region surrounding rs7307261 (KO/KO line, **Supplementary Fig. 3a**), as well as an edited line in which the A/A genotype was converted to G/G (G/G line, **Fig. 4a, Supplementary Fig. 3a**). We then differentiated these iPSC lines into microglial-like cells and validated their microglial identity by immunostaining for microglia-associated markers, including TMEM119, CX3CR1, and PU.1 (**Supplementary Fig. 3b**). We then used qPCR to examine *CD163* mRNA levels. We found no significant difference between A/A and edited G/G cells at baseline, suggesting that the rs7307261 A allele may not substantially affect *CD163* expression (**Fig. 4b**). However, ethanol exposure increased *CD163* mRNA levels in A/A microglia, whereas this increase was not observed in the knockout or edited G/G lines (**Fig. 4b**). Consistent with the mRNA expression results, immunostaining showed increased CD163 protein expression following ethanol exposure in the A/A line but not in the knockout or edited G/G lines (**Fig. 4c, d**). Together, these findings suggest that the rs7307261 A allele may support an ethanol-induced CD163 expression in human microglia.

**Figure 4.**
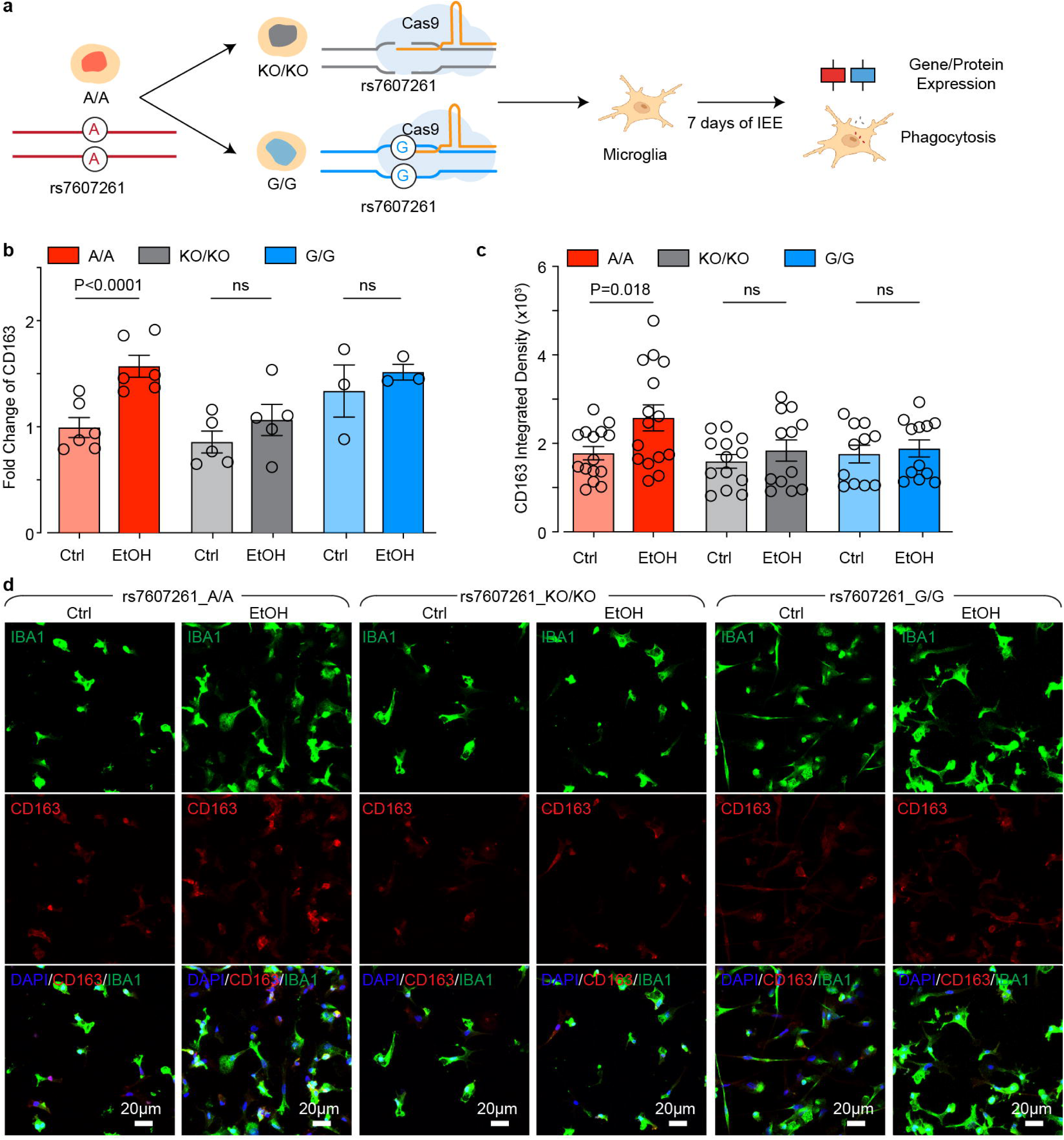
The rs7307261 A allele regulates ethanol-induced *CD163* expression. **(a)** Schematic of CRISPR/Cas9 editing of rs7307261 A/A iPSCs to generate KO/KO and G/G isogenic lines, followed by differentiation into microglia, 7 days of ethanol exposure, and assessment of gene/protein expression and phagocytosis. **(b)** Bar graph showing *CD163* mRNA expression measured by RT-qPCR in rs7307261 A/A, KO, and G/G. iPSC-derived microglial cells, with or without ethanol exposure. n = 6, 5, and 3 independent differentiation batches for A/A, KO/KO, and G/G respectively. Paired data were analyzed using a mixed-effect model with the Geisser-Greenhouse correction, followed by Šídák’s multiple-comparisons test. Data are presented as mean ± SEM. NS, not significant. **(c)** Bar graph showing quantification of CD163 immunostaining intensity in rs7307261 A/A (n = 15 images), KO/KO (n = 13 images), and G/G (n = 12 images) microglial cells with or without ethanol exposure. Statistical significance was assessed using ordinary one-way ANOVA followed by Šídák’s multiple-comparisons test. Data are presented as mean ± SEM. NS, not significant. **(d)** Representative immunofluorescence images of iPSC-derived microglial cells with A/A, KO/KO, and G/G genotypes under control and ethanol-treated conditions. CD163 is shown in red, IBA1 in green, and DAPI in blue.

### The rs7307261 A allele is associated with ethanol-induced microglial phagocytosis

Because CD163 has been linked to phagocytic responses in macrophages and microglia, we next examined whether rs7307261 genotype was also associated with phagocytic activity in microglial-like cells after ethanol exposure. The same CRISPR editing lines (i.e., rs7307261 A/A, KO/KO, and edited G/G microglial-like cells) were used for the phagocytosis assay (**Fig. 5a**).

**Figure 5.**
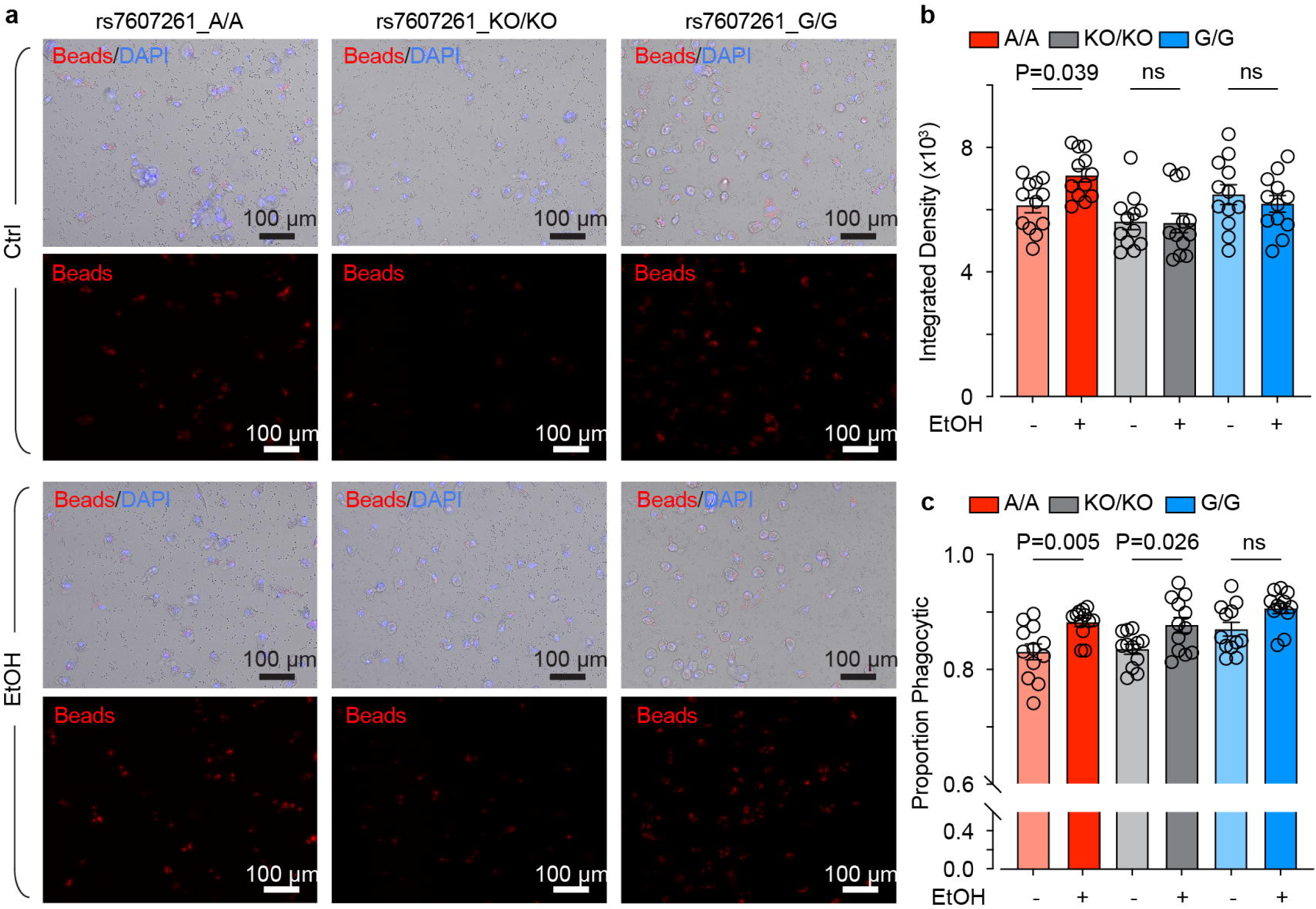
The rs7307261 A allele regulates ethanol-induced microglial phagocytosis. **(a)** Representative images of pHrodo-labeled zymosan bead uptake in A/A, KO/KO, and G/G microglial cells under control and ethanol-treated conditions. Beads are shown in red, and nuclei are labeled in blue. **(b)** Quantification of bead uptake by integrated density in A/A, KO/KO, and G/G microglial cells with or without ethanol exposure, n = 12 images from 3 independent batches per group. Statistical significance was assessed using ordinary one-way ANOVA followed by Šídák’s multiple-comparisons test. Data are presented as mean ± SEM. NS, not significant. **(c)** Quantification of the proportion of phagocytic microglia in A/A, KO/KO, and G/G microglial cells with or without ethanol exposure, n = 12 images from 3 independent batches per group. Statistical significance was assessed using ordinary one-way ANOVA followed by Šídák’s multiple-comparisons test. Data are presented as mean ± SEM. NS, not significant.

After intermittent ethanol exposure, A/A microglial-like cells showed a significant increase in overall phagocytic activity, whereas knockout and edited G/G cells did not show a significant change (**Fig. 5b**). We also observed an increase in the proportion of phagocytic cells in both A/A and KO/KO lines, but not in edited G/G lines (**Fig. 5c**). These results suggest that the rs7307261 variant is associated with altered microglial phagocytic activity after ethanol exposure.

## Discussion

AUD-related GWAS have identified risk variants and prioritized candidate genes using bulk RNA sequencing and chromatin interaction data^2^; however, these findings lack cell-type specificity and do not establish allele-specific regulatory effects or link the direction of these effects to gene expression changes or cellular function in AUD-relevant cell types. In this study, we addressed this gap by integrating microglia-specific ASoC mapping with AUD GWAS and AUD-related transcriptomic datasets. This approach prioritized rs7307261 A allele as an AUD risk allele associated with increased *CD163* expression under AUD- and ethanol-related conditions. Using human microglial-like cells, we further showed that rs7307261 A allele is associated with ethanol-induced upregulation of *CD163* and enhanced phagocytic activity, potentially through altered transcription factor regulatory networks. Together, these findings suggest that rs7307261 A allele may contribute to AUD-relevant microglial functional changes by regulating *CD163*-associated immune and phagocytic responses.

A key strength of our framework is the integration of multiple lines of evidence to prioritize AUD-associated variants and genes in microglia, followed by functional validation using multiple assays. Rather than relying on overlaps among genetic and functional annotations alone, our framework incorporates cell-type- and allele-specific chromatin accessibility and requires concordance in the direction of genetic risk, regulatory effects, and gene expression changes. We evaluated these relationships across complementary systems, including postmortem microglia from individuals with and without AUD and human iPSC-derived microglial-like cells with and without ethanol exposure. The convergence of genetic, epigenomic, and transcriptomic evidence across these datasets provides complementary support for regulatory variant-gene relationships relevant to AUD. The AUD risk allele (A allele) at rs7307261 showed allele-specific regulatory effects directionally consistent with increased *CD163* expression, while the surrounding chromatin region showed greater accessibility in AUD microglia. Moreover, the association between the A allele and increased *CD163* expression was observed in both postmortem microglia and iPSC-derived microglial-like cells under alcohol-related conditions. Thus, the rs7307261 and *CD163* relationship was supported by concordant genetic, chromatin, and gene expression evidence across complementary datasets and further strengthened by functional validation using CRISPR-mediated genome editing and cellular assays.

Although rs7307261 does not reach genome-wide significance in the AUD GWAS, this is expected given the highly polygenic nature of AUD, with genetic risk distributed across many variants with individually small effects. AUD is substantially heterogeneous, encompassing diverse symptoms and clinical features, and genetic variants may contribute differentially to specific aspects of the disorder. Consequently, variants associated with particular AUD-related features may show relatively modest associations with the broadly defined AUD phenotype.

Consistent with this possibility, GWASATLAS (https://atlas.ctglab.nl/PheWAS) shows that this variant is nominally associated with multiple cognitive, neurological, and psychiatric traits. A recent study^19^ further showed that GWAS significance rankings do not directly reflect biological effect sizes and are influenced by trait specificity and allele frequency. Thus, the absence of genome-wide significance alone does not preclude a functional role for rs7307261, particularly given the convergent genetic and functional evidence identified in our study.

Importantly, *CD163* itself also shows nominal associations with alcohol-related traits, including alcohol dependence (P = 0.005)^20^, beer/cider intake (P = 0.004) ^21^, and drinks per day (P = 0.043)^22^. *CD163* is a macrophage/microglial scavenger receptor ^23^. Although some studies indicate that in the normal brain, CD163 is expressed at low levels in microglia but is highly expressed in macrophages^24^, other studies show that microglia have increased CD163 expression under inflammatory or injury conditions, including stroke^25^, Alzheimer’s disease^26^, Parkinson’s disease^27^, Multiple Sclerosis^28^, and neuroinflammatory schizophrenia^29^. Functionally, CD163 is not only a hemoglobin-haptoglobin scavenger receptor but also exists in a soluble form that binds and neutralizes the cytokine TWEAK, preventing its interaction with the Fn14 receptor and suppressing downstream NF-κB and Notch signaling^30, 31^. This contributes to an overall anti-inflammatory phenotype. At the same time, CD163-high microglia/macrophages often retain or enhance their phagocytic capacity for debris and dead cells. Consistent with our results, increased *CD163* expression has been linked to enhanced microglial phagocytosis, as shown by Ferreira et al., who demonstrated that *CD163* upregulation promotes phagocytic activity in a neurodegenerative disease model^27^. More recently, CD163 has also been identified as a marker of interferon-activated microglia^32^, a subtype that regulates cortical development and behavior through neuronal phagocytosis^33^.

We found that chromatin accessibility at the rs7307261 A allele was increased in microglia from individuals with AUD compared with controls, and ethanol exposure increased *CD163* expression in A/A cells but not in edited G/G cells. These findings suggest that the regulatory effect of the rs7307261 A allele on *CD163* expression may depend on ethanol exposure, highlighting the importance of investigating gene regulation under disease-relevant conditions^34^. This is particularly relevant to AUD, as chronic alcohol consumption not only contributes to AUD-related symptoms but also increases risks for >200 diseases^35^. Understanding how genetic variants influence cellular responses to chronic alcohol consumption may therefore provide insights into individual differences in susceptibility to alcohol-related consequences and potentially inform future strategies for AUD management and recovery. Furthermore, the ethanol-dependent regulatory effect of rs7307261 may help explain its modest association with AUD in GWAS, as conventional case-control analyses may not fully capture genetic effects that vary with chronic alcohol consumption. A potential mechanism underlying this allele-specific regulation involves ETS-family transcription factors. Motif analysis predicted greater binding potential for ETS-family members, including FLI1, ELF2, ELK4, ELK1, and ETS1, at the rs7307261 A allele than at the G allele. These observations suggest a possible mechanism that allele-dependent differences in ETS transcription factor binding may indirectly influence ethanol-induced *CD163* expression, which remains to be experimentally validated.

Among numerus AUD-associated variants and genes, our integrative analyses prioritized rs7307261-*CD163* for detailed functional characterization. Several factors may account for this substantial reduction. First, recent large-scale single-nucleus transcriptomic studies of human postmortem brain tissue suggest that many AUD-associated molecular alterations occur in neuronal cells, whereas a much smaller subset is detected in microglia^36–38^. Our microglia-focused framework therefore captures only one component of the genetic architecture of AUD. Second, our prioritization framework was intentionally stringent, requiring convergence across multiple independent layers of evidence, including GWAS, microglial ASoC, eQTLs, DEGs, and chromatin accessibility, as well as concordance in the direction of genetic risk, allele-specific regulatory effects, and gene expression changes. Thus, variants that did not satisfy all of these criteria are not necessarily nonfunctional but could not be supported by the full set of evidence available in this study. Third, our experimental validation focused on responses to ethanol exposure in human microglial-like cells and therefore preferentially identifies variants whose regulatory effects are manifested under ethanol exposure. Other AUD-associated variants may act through different biological processes, cell types, developmental stages, or environmental conditions that are not captured by our experimental design.

This study has several limitations. First, although our ASoC, eQTLs, DEGs, and functional analyses suggest a potential regulatory relationship between rs7307261 and *CD163*, the molecular mechanism linking the A allele to altered *CD163* expression has not been experimentally established. In addition, the predicted allele-specific binding of ETS-family transcription factors at rs7307261 has not been directly demonstrated. Second, *CD163* is not specific to microglia but is also highly expressed in peripheral macrophages. Whether the regulatory effects observed in this study extend to peripheral macrophages remains to be determined. Third, the relatively modest sample sizes of both postmortem microglial and iPSC-derived microglial-like cell datasets limit statistical power, particularly for variants with small effects or low minor-allele frequencies. Larger datasets may therefore identify additional regulatory variant-gene relationships that the present study could not detect and further clarify the molecular mechanisms underlying the potential regulatory relationship between rs7307261 and *CD163*.

In summary, by integrating microglia-specific ASoC data with AUD GWAS, eQTLs, and DEGs, we identified the rs7307261 A allele as a functional AUD risk allele associated with ethanol-induced increased chromatin accessibility and C*D163* expression, potentially affecting microglial immune and phagocytic functions. These findings provide insights into the microglia-specific and stimulation-specific regulatory mechanisms underlying genetic susceptibility to AUD, highlighting the important role of microglia in alcohol-related neuropathology and providing a framework for linking genetic susceptibility to alcohol-related molecular and cellular phenotypes.

## Methods

### Integration of microglial ASoC and alcohol-related GWAS data

Microglia-specific ASoC regions were from Kozlova and colleagues^3^, who performed ATAC-seq in human iPSC-derived microglia derived from 38 subjects and identified variants with allele-specific chromatin accessibility. AUD-related GWAS summary statistics^2^ were downloaded from https://medicine.yale.edu/lab/gelernter/stats/ and the database of Genotypes and Phenotypes (phs001672). All GWAS results, including problem alcohol use and AUD-only (i.e., excluding those only having alcohol related problems but not diagnosed as AUD), were performed within each ancestry, cross-ancestry, and sex stratified^2^.

### Postmortem microglia snRNA-seq and snATAC-seq analyses

Single-nucleus RNA-seq and multiome (RNA + ATAC) data from postmortem caudate nucleus of individuals with AUD and controls were generated, processed, and annotated as previously described^11^. For each cell type, pseudobulk profiles were created by summing raw counts across all nuclei from each donor, and donors contributing fewer than 50 nuclei of that cell type were excluded. Peaks were called separately for each cell type using MACS2^39^ through the CallPeaks function in Signac^40^. This yielded microglial pseudobulk expression profiles (126 donors) and chromatin accessibility profiles (109,322 peaks; 70 donors) for donors with rs7307261 genotypes, obtained from nanopore long-read whole-genome sequencing.

Differential expression and accessibility were tested with DESeq2^41^ Wald tests on pseudobulk counts, with age and sex as covariates. Accessibility of the peak overlapping rs7307261 (chr12:7,970,288-7,971,265, GRCh38) was compared between AUD cases and controls using the design ∼ age + sex + AUD_status. To test the AUD effect within each genotype, donors were assigned to six groups based on rs7307261 genotype (A/A, G/A, G/G) and AUD status, and the design ∼ age + sex + group was fit, contrasting AUD cases with controls within each genotype for rs7307261 peak accessibility and for *CD163* expression, and A/A AUD cases with G/A AUD cases for *CD163* expression. Differences in CD163 expression between genotypes were tested with the design ∼ age + sex + genotype, both with and without AUD status as an additional covariate. For plotting, counts were normalized using DESeq2’s median-of-ratios size factors and plotted as ln(normalized count + 1). Coverage tracks were drawn with Signac’s CoveragePlot^40^.

### Microglia eQTL analysis

Genotypes were called from the snATAC-seq data of each donor as described previously^42^: per-donor BAM files were generated with sinto (https://timoast.github.io/sinto/), and variants were called with the Sentieon germline pipeline^43^ (Haplotyper and joint calling with DNAseq), followed by GATK Variant Quality Score Recalibration^44^. Cis-eQTL mapping was performed with tensorQTL^45^ using microglial pseudobulk expression of all genes and all variants located within microglial ATAC-seq peaks with minor allele frequency ≥ 0.05, within +-500 kb of each gene’s transcription start site. Sex, age, and AUD status were included as covariates. Nominal variant-gene associations were used for integration with the ASoC and GWAS data.

### Allele-specific accessibility in microglia from postmortem donors

Reads carrying each rs7307261 allele were counted in microglial nuclei from the snATAC-seq BAM files using VarTrix (v1.1.22, coverage scoring; https://github.com/10XGenomics/vartrix) and summed per donor.

### Human iPSC lines

Twenty-five human iPSC lines were generated from lymphocytes and lymphoblastoid cells of participants in the Collaborative Study on the Genetics of Alcoholism (COGA) (NIAAA/COGA Sharing Repository) (**Supplementary Table 4**). Cryopreserved lymphocytes were thawed, expanded as erythroblasts, and reprogrammed with Sendai virus (CytoTune, Life Technologies). Reprogramming and pluripotency assessment were performed by Sampled (Piscataway, NJ). We maintained iPSCs feeder-free on Matrigel (Corning, 354234) in mTeSR Plus medium (STEMCELL Technologies, 100-0276) and passaged them with Accutase (STEMCELL Technologies, 07920). All experiments were conducted in accordance with Rutgers Biosafety protocol #13-444.

### Generation of rs7307261-edited iPSC lines by CRISPR/Cas9

iPSC lines were electroporated with precomplexed Cas9 protein (Life Technologies) and gRNA (custom-designed using TrueDesign software, Life Technologies) using the Neon XT system (Life Technologies) with Alt-R Cas9 Electroporation Enhancer (IDT). The target sequence was 5’-TGTATGCGTaAGTCTTTGGG-3’, with the lowercase “a” indicating the position and identity of the targeted SNP. Cells were recovered in StemMACS iPS-Brew XF (Miltenyi) supplemented with the combination of chroman 1, emricasan, polyamines, and trans-ISRIB (CEPT)^46^. Cells were diluted into 96-well plates, followed by harvesting wells with single colonies for expansion. Clones were assayed by Sanger sequencing of a PCR product (primers: Forward 5’-TCCCTTTCTTGCACTCGTCT-3’ and Reverse 5’-GAGAAGACGCCGATTCCTGA-3’) as interpreted by ICE software (EditCo). For heterozygotes, we performed single-molecule sequencing on the PCR product (Plasmidsaurus, Inc.) and counted specific deletions on individual molecules. For clone F, counting 2,666 individual reads detected 47.3% and 45.3% of the 18 and 17 bp deletions, respectively.

### Microglia progenitor cell differentiation

Microglial-like cell differentiation was performed as previously described ^17^. iPSCs were induced to form yolk sac embryoid bodies (YS-EBs) in mTeSR Plus medium with BMP4 (50 ng/ml; PeproTech, #120-05ET), VEGF (50 ng/ml; PeproTech, #100-20A), and SCF (20 ng/ml; PeproTech, #300-07) for 6 days. YS-EBs were then plated in X-VIVO 15 medium (Lonza, #02-060Q) containing IL-3 (25 ng/ml; PeproTech, #200-03) and M-CSF (100 ng/ml; PeproTech, #200-25), generating primitive macrophage precursors (PMPs) that appeared after ∼4 weeks and were maintained for >6 months. For maturation, PMPs were cultured for 14 days in microglial-like cell differentiation medium consisting of DMEM/F12 (HyClone, #SH30023.01) supplemented with N-2 (Gibco, #17502048), IL-34 (100 ng/mL; PeproTech, #200-34), and GM-CSF (10 ng/mL; PeproTech, #300-03).

### Intermittent ethanol exposure

After a 7-day maturation period in microglia differentiation medium, human-iPSC-derived microglial cells were exposed to a 7-day chronic intermittent ethanol paradigm^17, 47, 48^. On the first day of treatment, we removed all medium from each well and replaced it with medium supplemented with ethanol (200 proof, non-denatured; Decon Laboratories, Cat# 2716) to achieve peak concentrations of 0 or 75 mM. Accounting for evaporation, this exposure paradigm produced a mean 24-hour ethanol concentration of 28.79 mM, corresponding to a blood alcohol concentration of approximately 0.13^48^. On days 2, 3, 5, and 6, half of the medium was removed from each well, ethanol was added to the removed medium at twice the target concentration, and the mixture was returned to the cultures. On days 4 and 7, we performed a half-medium change by replacing half of the existing medium with fresh medium containing ethanol at twice the target concentration. On day 8, cultures were collected for downstream analyses.

### RNA purification and RT-qPCR

Total RNA was isolated from ∼1.5–2 million cells using the RNeasy Plus Mini Kit (Qiagen, Cat# 74136) following the manufacturer’s protocol. Lysates were homogenized with a QIAshredder, genomic DNA removed with gDNA Eliminator columns, and RNA purified on RNeasy spin columns with RW1 and RPE washes. RNA was eluted in RNase-free water and quantified using a NanoDrop spectrophotometer (Thermo Scientific).

RT-qPCR was performed with PowerUp SYBR Green Master Mix (ThermoFisher, Cat# A25743) on a QuantStudio 3 system under the following cycling conditions: 50 °C for 2 min, 95 °C for 2 min, then 40 cycles of 95 °C for 15 s and 60 °C for 1 min. Primers were designed using NIH Primer-BLAST, and reactions were run in duplicate across five biological replicates.

HsaCD163qPCR_Ex13_14: Forward primer: CAAAAAGCCACAACAGGTCG. Reverse primer: CAACAGAACAACCCCAAGGA

### Immunocytochemistry

Cells were fixed in 4% PFA (PBS) for 15 min at room temperature and blocked with 5% goat serum, 4% BSA, and 0.05% Triton X-100 (PBS) for 60 min. Primary antibodies diluted in blocking buffer included chicken anti-IBA1 (Aves Labs, Cat# IBA10020, RRID: AB_2910556) and rabbit anti-CD163 (Abcam, Cat# ab182422, RRID: AB_2753196), mouse anti-TMEM119 (Cell Signaling Technology, catalog #41134S, RRID: AB_3094467), rabbit anti-PU.1 (Cell Signaling Technology, catalog #2266S, RRID: AB_10692379), rabbit anti-CX3CR1 (Bio-Rad, catalog #AHP1589, RRID: AB_2087421). Samples were incubated with primary antibodies for 1 h at room temperature or overnight at 4 °C. After five washes with PBST (0.05% Triton X-100 in PBS), secondary antibodies conjugated to Alexa Fluor 488 or 546 (anti-mouse, anti-rabbit, and/or anti-chicken; Thermo Fisher Scientific) were applied at 1:500 in blocking buffer and incubated for 1 h at room temperature. Following five additional PBST washes, coverslips were mounted with Fluoroshield containing DAPI (Sigma). Microscopic images were captured using a confocal microscope (Zeiss LSM700) for further examination and data acquisition.

### Phagocytosis assay

Human iPSC-derived microglia were seeded in 96-well plates following a 7-day intermittent ethanol exposure paradigm with 0 mM or 75 mM ethanol. On day 7 of ethanol exposure, Hoechst was diluted 1:1000 and added to the microglia to label nuclei, followed by incubation at 37°C for 10 minutes. Zymosan bioparticles (Thermo Fisher Scientific, catalog #P35364) were prepared by vortexing and ultrasonication for 5 minutes. The prepared zymosan bioparticles were then added to live microglia in a bath solution containing, in mM: 140 NaCl, 5 KCl, 10 HEPES, 2 CaCl₂, 2 MgCl₂, and 10 glucose, pH 7.4. Cells were incubated for 2 hours at 37°C. Live-cell images were acquired using a Keyence BZ-X800 microscope with BZ-X800 software, and ImageJ was used for data analysis.

## Statistical analysis

Statistical analyses were performed using GraphPad Prism v10.0 (GraphPad Software, La Jolla, CA). Data are presented as mean ± SEM unless otherwise stated. Analyses were based on biologically independent replicates, as described in the figure legends. To evaluate the impact of chronic ethanol exposure, results were normalized to baseline activity. Differences in ethanol effects across genotypes were assessed using one-way ANOVA followed by Fisher’s least significant difference post hoc test. Within-group comparisons were performed using paired t-tests before and after ethanol exposure. Statistical significance was defined as P < 0.05 (*), P < 0.01 (**), and P < 0.001 (***), with NS indicating no significant difference.

## Acknowledgements

The Collaborative Study on the Genetics of Alcoholism (COGA), Principal Investigators B. Porjesz, V. Hesselbrock, A. Agrawal; Scientific Director, A. Agrawal; Translational Director, D. Dick, includes nine different centers: University of Connecticut (V. Hesselbrock); Indiana University (H.J. Edenberg, T. Foroud, Y. Liu, M.H. Plawecki); University of Iowa Carver College of Medicine (A. Andersen S. Kuperman); SUNY Downstate Health Sciences University (B. Porjesz, J. Meyers); Washington University in St. Louis (L. Bierut, A. Agrawal, S. Hartz); University of California at San Diego (M. Schuckit); Rutgers University (D. Dick, R. Hart, J. Salvatore, J. Tischfield); The Children’s Hospital of Philadelphia, University of Pennsylvania (L. Almasy); Icahn School of Medicine at Mount Sinai (A. Goate, P. Slesinger); and Howard University (D. Scott). Other COGA collaborators include: M. Hesselbrock, K. Manning (University of Connecticut); D. Lai, J. Nurnberger Jr., L. Wetherill, A. Miller, X. Xuei, (Indiana University); J. Kramer (University of Iowa), G. Chan (University of Iowa; University of Connecticut); C. Kamarajan, A. Pandey, D.B. Chorlian, P. Barr, S. Kinreich, G. Pandey, Z. Neale, C. Chatzinakos, J. Zhang, S. Saenz deViteri, A. Bingly (SUNY Downstate); G. Pathak (Icahn School of Medicine at Mount Sinai); A. Anokhin, K. Bucholz, F. Dong, A. Hatoum, E. Johnson, J. Rice, S. Saccone (Washington University); F. Aliev, Z. Pang, S. Kuo, S. Brislin, (Rutgers University). We continue to be inspired by our memories of Henri Begleiter and Theodore Reich, founding PI and Co-PI of COGA, and also owe a debt of gratitude to other past organizers of COGA, including Ting-Kai Li, P. Michael Conneally, Raymond Crowe, and Wendy Reich, for their critical contributions. This national collaborative study is supported by NIH Grant U10AA008401 from the National Institute on Alcohol Abuse and Alcoholism (NIAAA) and the National Institute on Drug Abuse (NIDA).

Y. Liu and D. Lai are supported by National Institutes of Health award number AA031176.

X. Li, R. Hart, and P. Pang are supported by National Institutes of Health award number R01AA023797.

## Conflict of Interest

The authors declare no conflict of interest.

## Author contributions

Z.P.P., D.L., and R.P.H. conceived and supervised the study. X.L., J.S.M., S.G., H.D.N., T.S., and W.Y. performed the experiments. X.L. and N.C.G. analyzed and interpreted the data, prepared the figures, and drafted the manuscript. J.D. and Y.L. contributed resources, data interpretation, and manuscript editing. All authors reviewed and approved the final manuscript.

## Data Availability

The postmortem human brain data analyzed in this study are available in the Gene Expression Omnibus (GEO, GSE277313). The raw RNA-seq data from human microglial cells are available in GEO (GSE255988). All other data that support the findings of this study are listed in Supplementary Data. Source data are provided with this paper.

## Code Availability

No custom computational packages or extensive computer code were developed in this study. However, R scripts used to utilize existing packages are available from the corresponding author upon request.

---

**Supplementary Figure 1. The potential ASoC-AUD variant-gene pairs identified in human microglial cultures and postmortem microglia. (a)** Manhattan plot showing the distribution of microglial ASoC-AUD variants with their alleles tested. **(b)** Manhattan plot showing the chromosomal distribution of ASoC-AUD risk variant-gene pairs associated with the overlapping upregulated DEGs. Dot size represents the eQTL slope, indicating the magnitude of the regulatory effect. **(c)** Manhattan plot showing the chromosomal distribution of ASoC-AUD risk variant-gene pairs associated with the overlapping downregulated DEGs. Dot size represents the eQTL slope, indicating the magnitude of the regulatory effect.

**Supplementary Figure 2. Chromatin accessibility at rs7307261, predicted TF binding motifs, and potential target gene expression. (a)** Chromatin accessibility at the rs7307261 region in AUD and control postmortem microglia. The red dashed line labels the genomic position of rs7307261. **(b)** JASPAR 2026 motif analysis of the genomic sequence spanning ±10 bp around rs7307261. Sequence logos show predicted FLI1 and ELF2 binding motifs overlapping the rs7307261 locus, with the A allele positioned within the motif core and highlighted in pink.

**Supplementary Figure 3. Generation and validation of rs7307261-edited iPSC-derived microglial-like cells. (a)** Sanger sequencing validation of CRISPR/Cas9-edited rs7307261 iPSC lines. The parental rs7307261 A/A line underwent the same CRISPR/Cas9 editing procedure and served as an isogenic control(top). The rs7307261_KO/KO line contains a 17-bp/18-bp deletion surrounding the rs7307261 locus (middle), while the edited rs7307261_G/G line shows conversion of the rs7307261 allele to G/G (botton). **(b)** Representative immunofluorescence images of iPSC-derived microglial-like cells carrying rs7307261_A/A, rs7307261_KO/KO, and edited rs7307261_G/G genotypes. Cells were stained for microglial markers TMEM119, CXCR3, PU.1, and IBA1, with DAPI labeling nuclei. Scale bar, 50 μm.

## Notes

### Competing Interest Statement

The authors have declared no competing interest.

