## Supplementary figures and images for "Allele-specific open chromatin analysis revealed allelic bias of alcohol-induced gene expression in human microglia"

### supplemetal figure1

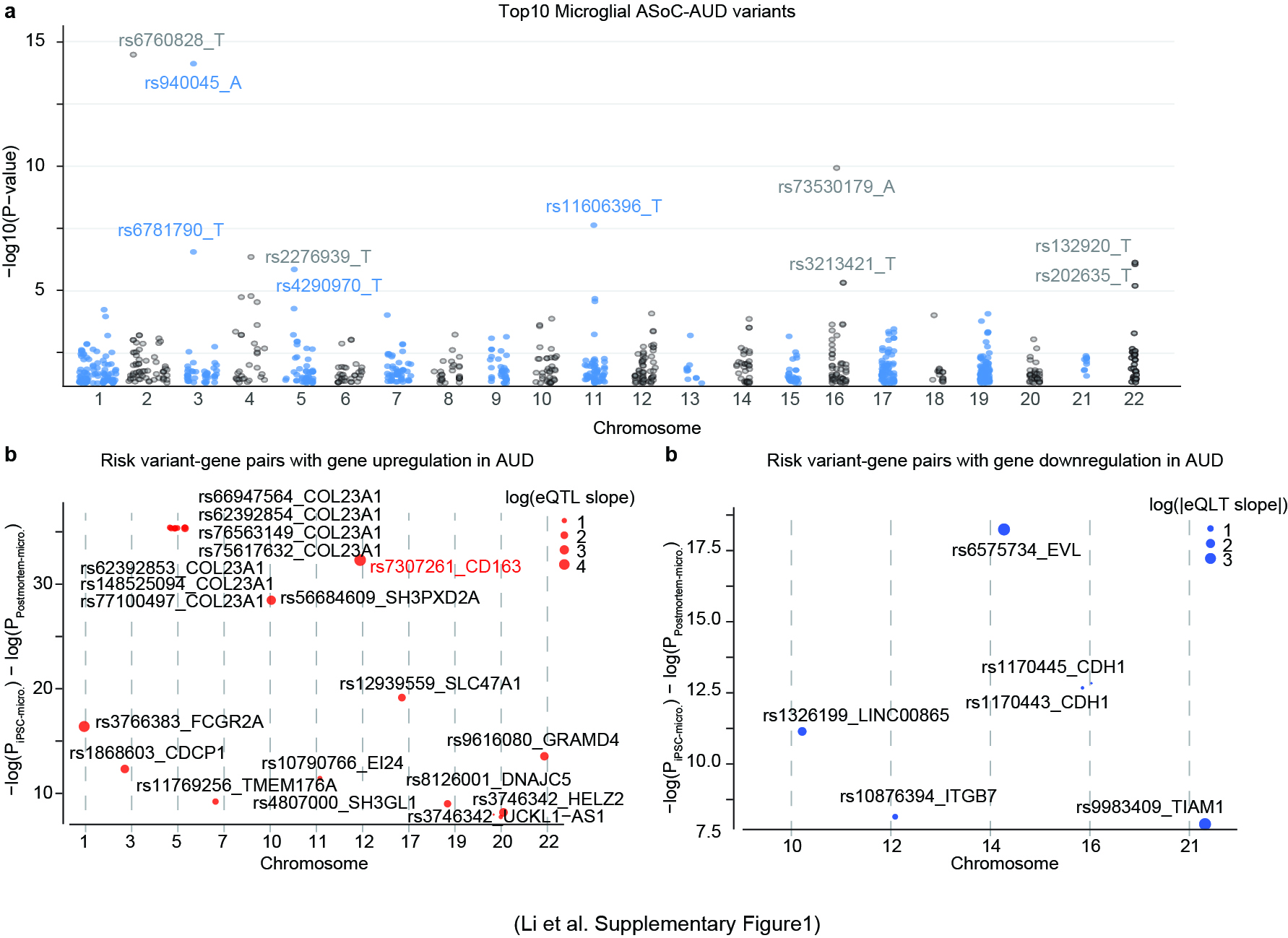

### supplemetal figure2

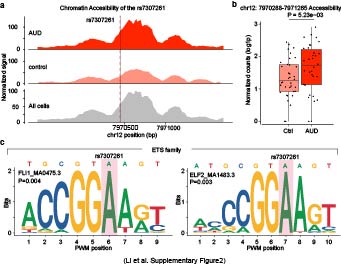

### supplemetal figure3

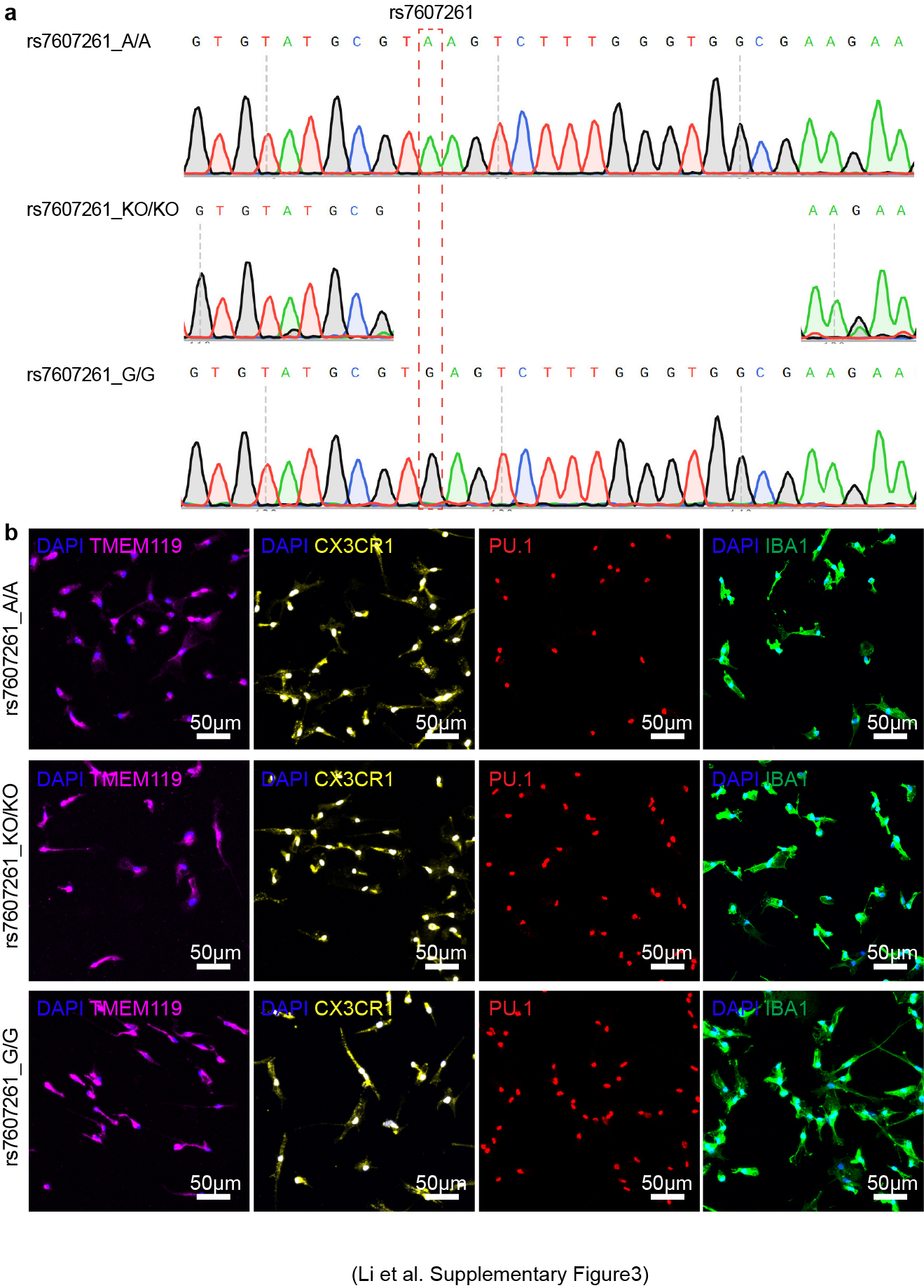
